# Parasitoids reduce growth rate in oak-feeding caterpillars

**DOI:** 10.1101/2025.05.23.655719

**Authors:** Freerk Molleman, Ahmet Tambay, Soumen Mallick, Stephanie Llopis, Andreas Prinzing, Urszula Walczak

## Abstract

1. Koinobiont parasitoids have been shown to both reduce and increase the growth rate of caterpillars. However, no comprehensive study has been conducted on all caterpillar hosts of a given plant species. Moreover, most of the existing case studies are on large caterpillars attacking crop plants. This may not be representative of the effect of parasitoids on caterpillar growth on trees.
2. We measured the growth rate of parasitised and non-parasitised spring caterpillars collected from oak trees (*Quercus robur & Q. petraea*) in France and Poland.
3. Across six species of Lepidoptera, parasitoids reduced caterpillar growth rate on average.
4. Further studies should determine if the reduced growth rate is accompanied by lower consumption rates, so that the plant directly benefits from attracting parasitoids when they are attacked by caterpillars.

## INTRODUCTION

Parasitoids typically kill a significant number of herbivorous insects such as caterpillars of Lepidoptera (Price et al., 1980). Some of these parasitoids paralyse their host permanently (idiobionts), but most paralyse their host only temporarily (koinobionts), so that the host can continue feeding and the parasitoids only prevent reproduction of their host (Mackauer & Sequeira, 1993). Therefore, parasitoids may be regarded as of less immediate benefit to individual plants than predators that kill herbivorous insects immediately. Plants attracting parasitoids (e. g. by emitting herbivore-induced plant volatiles; Cuny & Poelman, 2022) may die before profiting from the delayed defense offered by parasitoids. Moreover, neighbouring conspecific plants would equally profit, irrespective of whether they attracted parasitoids or not. Overall, whether plants are under selection to attract parasitoids will depend a lot on how parasitoids affect the growth of the caterpillars.

Counter-intuitively, parasitoids can both decrease and increase the growth of their hosts. Parasitoids can reduce caterpillar growth rate as a passive disease response or as active manipulation by the parasitoid, but may also induce caterpillars to grow for a longer time and become larger (reviewed in Cuny & Poelman, 2022). To manipulate host growth, the adult parasitoid can inject factors along with the eggs into the host, such as endogenous viruses, venom, and teratocytes (Cuny & Poelman, 2022). Parasitoid larvae can also produce factors that affect host growth (e.g. Brown et al., 1993). Reduced growth rate may be due to the parasitoid reducing the feeding rate of its host (e.g. Adamo et al., 2016). Reduced growth rate is generally accompanied by reduced leaf consumption rate, but in the laboratory, overall consumption may not be affected, or can even be increased if parasitoids prolong the host’s development time (e.g. Alleyne & Beckage, 1997; Moore et al., 2020). According to a recent review (Cuny & Poelman 2022), a similar number of cases has been described in which parasitoids increase or decrease growth (often quantified as maximum size rather than growth rate) of their host insects. However, no comprehensive study has been conducted on the community of caterpillar hosts of a given plant species. Moreover, most of these case studies are on large caterpillars attacking crop plants. This may not be representative of the effect of parasitoids on caterpillar growth on trees.

To test if parasitoids reduce the average growth rate of caterpillars, we compared the growth rate of non-parasitised caterpillars with that of parasitised caterpillars collected from oak trees (*Quercus petraea* (Matt.) Liebl. & *Q. robur* L.) in two forests.

## METHODS

### Overview

We collected and reared spring-feeding caterpillars from the canopy of mature oak trees (*Q. petraea, Q. robur*) in the Forêt de Rennes in Brittany, western France (2012 and 2019), and the Puszcza Zielonka forest in western Poland (2021 and 2022).

### Caterpillar rearing in France 2012

We sampled branches from 19 oak trees (*Q. petraea, Q. robur*) shortly after budburst (11^th^ April to 11^th^ May 2012). We transported the branches in plastic bags to the lab and reared the associated caterpillars in Petri dishes at room temperature using locally obtained *Q. robur* leaves. We measured the length of the caterpillars using callipers every alternate day (24^th^ April to 24^th^ May). We assigned the caterpillars to surrogate species and linked these later to species using individuals that we reared to adulthood and identified. We noted when a caterpillar produced a parasitoid or pupated.

### Caterpillar rearing in France 2019

We collected branches from 24 oak trees (*Q. petraea, Q. robur*) in spring (26^th^ March to 23^rd^ May 2019) and reared the caterpillars in the lab We reared all caterpillars separately in Petri dishes in a climate room mimicking the average diurnal change in temperature, daylight, and relative humidity of the study forest. We fed the caterpillars with freshly emerged leaves. In addition to classifying caterpillars into surrogate species and linking these to reared adults, we used DNA barcoding to provisionally identify caterpillars that did not reach adulthood. The caterpillars were weighed on a semi-microbalance on average twice a week.

### Caterpillar rearing in Poland 2021

We collected branches from the canopy of four sessile oak trees (*Q. petraea)* shortly after budburst (May 14^th^ 2021). We cut the branches into pieces and transported them to the laboratory in plastic bags. The next day, we searched the branches for caterpillars and placed each individually in a 50 ml cup. *Operopthera brumata* (L.) (Geometridae) caterpillars could be putatively recognised, and other morphologies were given surrogate-species names. We reared the caterpillars individually on leaves from these trees in a climate chamber with a constant temperature of 20°C and relative humidity of 75%. Three days later, we removed the frass and placed a new leaf inside each cup. We weighed the caterpillars the day after collection before adding food, and six days later, as they were transferred to a new cup. Thus, on both occasions, caterpillars could not feed for a few hours before weighing. Caterpillars were henceforth fed locally obtained oak leaves, and we noted death, parasitoid exit, pupation, and eclosion dates. We identified all eclosed Lepidoptera to species using dissection of genitalia when needed (Lepiforum, 2020; Razowski, 1983, 1987; Wheeler, 2023) to then link caterpillar surrogate species to species.

### Caterpillar rearing in Poland 2022

The study in 2022 was similar to that in 2021, but sampling ten trees (six on May 11^th^ and four on May 16^th^), and their mass change was measured over a five-day period (described in more). Unfortunately for the current purpose, this time we avoided rearing caterpillars that appeared parasitised.

### Data analysis

To estimate caterpillar mass from their length (data from France 2012), we assumed that caterpillars had a cylindrical shape and thus utilised the formula ‘π × (height / 2)^3^ × length’ to calculate their volume. For each species, we inferred height from length by determining the height-length ratio from lateral photographs of caterpillars. We thus estimated that the height-length ratios were 0.1563 for *Cosmia trapezina* (L.) (Noctuidae), 0.1364 for *O. brumata*, and 0.1507 for *Orthosia cruda* (Den. & Schiff.) (Noctuidae). We finally assumed that the mass density of caterpillars is similar to water, to translate estimated volume into estimated body mass.

Quantifying caterpillar growth rate is complicated because caterpillar growth occurs in phases, and mass gain changes with body size. Growth pauses during moulting (change of instar), and caterpillars lose considerable body mass while preparing to pupate (Tammaru & Esperk, 2007). Growth rate can be approximated as ‘(final mass-initial mass)^1/3^ / development time’ (including the moulting, Tammaru & Esperk, 2007; Supplement 1). We visualised the growth curves of oak-feeding caterpillars in our study for individuals with at least three size measurements (France 2012 and 2019; Supplement 1). To calculate growth rate, we took the first measurement as the initial mass. In an attempt to avoid the mass loss associated with pupation, when multiple measurements were available for an individual (2012 and 2019), we selected as final mass the measurement before mass gain decreased (Supplement 1). For 2012, we also excluded caterpillar measurements that were longer than 16 mm. When we had only two measurements (Poland 2021, 2022), we used a short period (5 □ 6 days) over which to calculate growth rate. We finally excluded caterpillars that lost rather than gained mass from analysis. These may have been pupating or unhealthy, and possibly included instances of mass loss induced by parasitoids. We also excluded the few instances of dipteran parasitoids.

We compared the growth rate of non-parasitised caterpillars that reached the pupal stage with those that were recorded as parasitised. We included only cases where, during a given year, we measured the growth rate of at least one parasitised and one non-parasitised individual for the species, and only species that had multiple observations in both categories across years. To account for methodological differences between years and differences between species, we used generalised estimating equations (GEE) with the interaction between year and species as the id, implemented using the *geepack* package in R (Højsgaard et al., 2024; R_Core_Team, 2025). This is similar to using random effects in mixed models, but allows for a wide range of sample sizes, with many categories containing only one observation (Liang & Zeger, 1986).

## RESULTS

### Overview

We analysed the growth rate of 246 non-parasitised and 34 parasitised caterpillars belonging to five known species and one unidentified surrogate species. These were *Acrobasis repandana* (Fabricius, 1798) (Pyralidae), *C. trapezina, Eudemis profundana* (Denis & Schiffermuller, 1775) (Tortricidae), *Op. brumata, Or. cruda*, and an unidentified leaf roller. The parasitoids were koinobiont Hymenoptera. The caterpillar growth rates we calculated varied considerably within species, both for non-parasitised and parasitised caterpillars (Figure 1). Overall, parasitism reduced caterpillar growth rate (Table 1). *O. brumata*, often one of the main herbivores on these oak trees, was the most common species in our data, and showed a significant reduction in larval growth rate (Figure 1; effect estimate = -0.0057, p < 0.001). Of the 62 caterpillars that lost mass (excluded from the above analyses), 12 were recorded as parasitised.

**Table 1.** Results of the Generalised Estimating Equations model for the effect of parasitation on caterpillar growth rate with family set as Gaussian, id as the interaction between species and year, the correlation structure as exchangeable, and link as identity. The number of clusters was 100, with a maximum cluster size of 60.

|  | Estimate | S. e. | Wald | P-value |
| --- | --- | --- | --- | --- |
| Intercept | 0.028 | 0.0016 | 312.3 | < 0.001 |
| Parasitation | -0.004 | 0.0016 | 7.6 | **0.006** |
| Estimated Scale Parameters: |  |  |  |  |
|  | Estimate | S. e. |  |  |
| Intercept | 0.00027 | < 0.001 |  |  |
| Estimated Correlation Parameters: |  |  |  |  |
|  | Estimate | S. e. |  |  |
| Alpha | 0.787 | 0.0842 |  |  |

**Figure 1.**
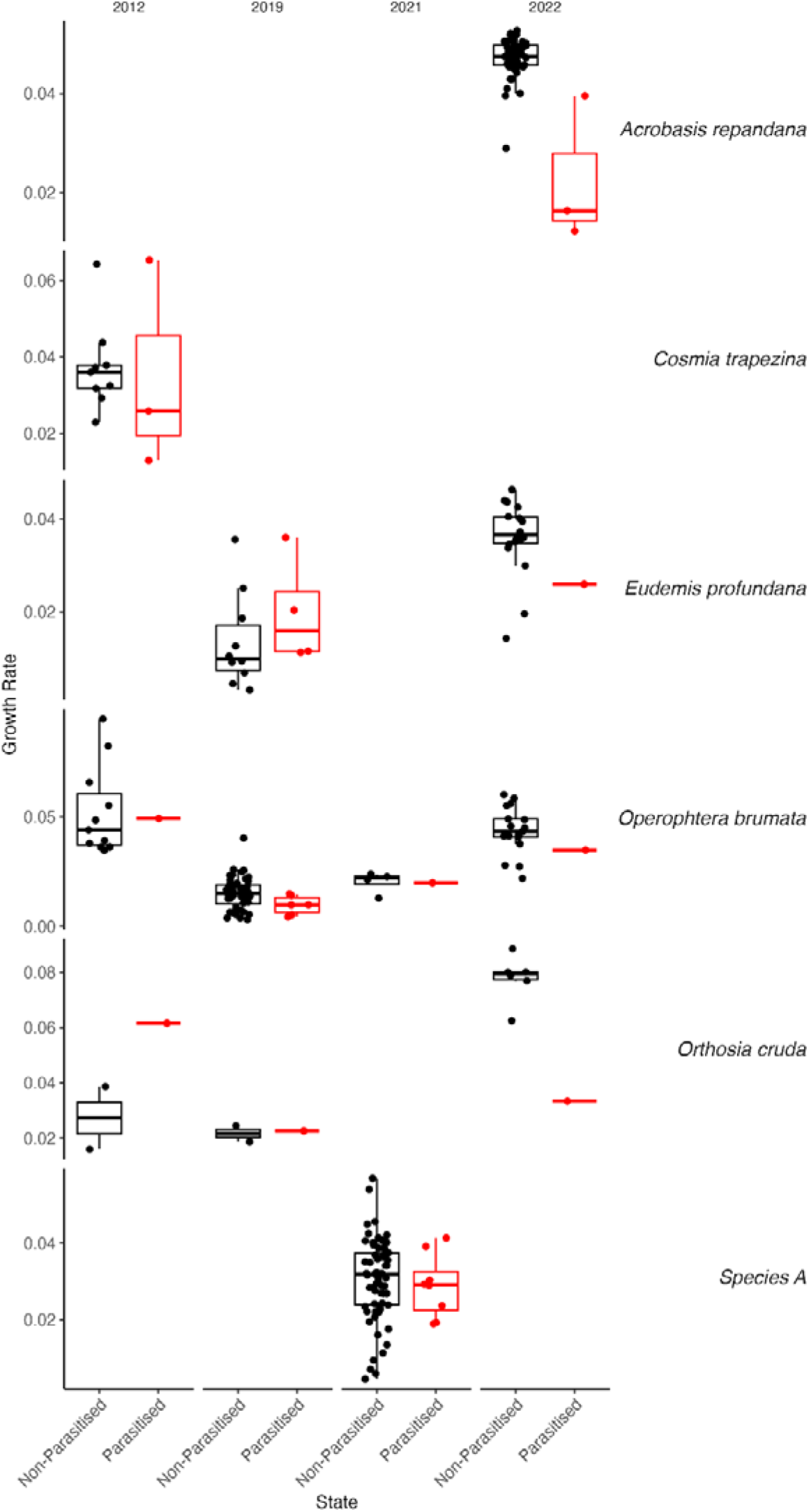
Scatter plot and boxplots of growth rate (mass increase (mg)^1/3^ / time interval (days)) of non-parasitised (in black) and parasitised (in red) caterpillars reared during four springs. The central line within each box represents the median, the box boundaries denote the interquartile range, and the whiskers extend to the smallest and largest values within 1.5 times the interquartile range. Data points beyond this range are displayed as outliers. Boxplots for species with few data points need to be interpreted with caution.

## DISCUSSION

We measured the growth rate of non-parasitised and parasitised spring caterpillars collected from oak trees (*Q. robur & Q. petraea*) in France and Poland, and found that across six species of Lepidoptera, parasitised caterpillars had a lower growth rate. However, some parasitised caterpillars showed growth rates as high or higher than the average of non-parasitised ones. Notably, by excluding caterpillars that lost rather than gained mass, and in 2022 excluding caterpillars that appeared parasitised, we probably excluded some cases in which parasitoids severely reduced caterpillar growth rate, so that our analysis likely underestimates the average reduction in caterpillar growth rate.

While our study estimated an overall effect of parasitoids on caterpillar growth rate, the parasitoids were not identified beyond order, so that we cannot compare the effects of growth rate that various clades of parasitoids may have, or e.g. correlations with the degree of host specialisation. With identified parasitoids, we could also ask whether certain caterpillar traits (e.g. host-plant specialisation, phenology, size, abundance) make them more or less prone to growth rate reduction. It is also important to note that our methods could not fully avoid noise in the data due to the pauses in growth during moulting and mass loss before pupation, as well as reduced growth rates that may be due to pathogens, or even parasitoids that developed inside caterpillars but never exited from them. Moreover, some of the caterpillars were shelter builders (*A. repandana, E. profundana*, and species A), so by breaking their shelters when we weighed or fed them, their growth rates were probably reduced (LoPresti & Morse, 2013). This extra stress might, for example, exacerbate the effect of parasitoids on growth rate. A further source of noise may be the stomach content, which may have varied considerably among the caterpillars in France that were removed from their food right before weighing. In Poland, caterpillars had been kept without food for several hours before weighing so that their digestive systems were probably empty, reducing this source of noise. Measuring caterpillar length may avoid this source of noise, but has its own limitations due to caterpillar stretchability and changing allometry during ontogeny. Finally, further noise was probably introduced when attempting to identify caterpillars that died based on surrogate species or DNA barcoding To get more accurate data on how parasitoids affect larval growth rate, we need controlled experiments in which caterpillars are weighed frequently, their development is monitored, and caterpillars are infected with known parasitoid species (e.g. Moore et al., 2020), but this is hardly feasible for a diverse caterpillar-parasitoid community associated with a species of tree. Nevertheless, overall, we have shown that parasitoid wasps reduce caterpillar growth rate across a wide variety of caterpillar and parasitoid species associated with oak trees in northern Europe.

Further studies need to address to what extent the reduced growth rates of parasitised caterpillars result in reduced leaf consumption. If slower-growing caterpillars indeed consume less leaf material, parasitoids would immediately benefit the individual plant, albeit this benefit being smaller than that of attracting predators that kill caterpillars immediately. Even if slower-growing caterpillars consume more by increasing development time, in the field, the slower growth rate will expose them to other natural enemies and the elements for a longer period. Moreover, as leaves become more defended as they develop (Feeny, 1970), slower growth will lead to spring caterpillars experiencing worse food quality, further reducing their growth and survival rate. So, reduced growth rate with extended development could still reduce the overall leaf damage under field conditions. Notably, the time spent feeding also affects mortality as caterpillars are more likely to be attacked during feeding (Bernays, 1997), but we have little insight into how parasitoids affect caterpillar feeding behaviour (see e.g. Adamo et al., 2016; Miles et al., 2023). Therefore, to establish the link between parasitoids manipulating host growth and the resulting leaf consumption, we need both matching data on leaf consumption and growth (e.g. Moore et al., 2020), as well as caterpillar mortality in the field (e.g. Chen et al., 2017; Remmel et al., 2011). If indeed reduced growth rate is accompanied by reduced leaf consumption in the field, our results would suggest that individual trees have an immediate benefit from recruiting parasitoids, and would thus be selected to do so.

## Supporting information

Appendix 1 growth curves

## Supplements

Supplement 1: Growth curves of caterpillars.

## Notes

### Competing Interest Statement

The authors have declared no competing interest.

### Funding

- NCN: 2018/29/B/NZ8/00112

## REFERENCES

Adamo, S.A., Kovalko, I., Turnbull, K.F., Easy, R.H., & Miles, C.I. (2016) The parasitic wasp Cotesia congregata uses multiple mechanisms to control host (Manduca sexta) behaviour. Journal of Experimental Biology, 219, 3750–3758.

Alleyne, M. & Beckage, N.E. (1997) Parasitism-induced effects on host growth and metabolic efficiency in tobacco hornworm larvae parasitized by Cotesia congregata. Journal of Insect Physiology, 43, 407–424.

Bernays, E. (1997) Feeding by lepidopteran larvae is dangerous. Ecological Entomology, 22, 121–123.

Brown, J.J., Kiuchi, M., Kainoh, Y., & Takeda, S. (1993) In vitro release of ecdysteroids by an endoparasitoid, Ascogaster reticulatus Watanabe. Journal of insect physiology, 39, 229–234.

Chen, W.-B., Vasseur, L., You, M.-S., Li, J.-Y., Wang, C.-X., Meng, R.-X., & Gurr, G.M. (2017) Parasitised caterpillars suffer reduced predation: potential implications for intra-guild predation. Scientific Reports, 7, 42636.

Cuny, M.A. & Poelman, E.H. (2022) Evolution of koinobiont parasitoid host regulation and consequences for indirect plant defence. Evolutionary Ecology, 36, 299–319.

Feeny, P.P. (1970) Seasonal changes in oak leaf tannins and nutrients as a cause of spring feeding by winter moth caterpillars. Ecology, 51, 565–581.

Højsgaard, S., Halekoh, U., Yan, J., & Ekstrøm, C. (2024) Package ‘geepack’. R package version, 1–2.

Lepiforum, e.V. (2020) Bestimmungshilfe für die in Europa nachgewiesenen Schmetterlingsarten.

Liang, K.-Y. & Zeger, S.L. (1986) Longitudinal data analysis using generalized linear models. Biometrika, 73, 13–22.

LoPresti, E. & Morse, D. (2013) Costly leaf shelters protect moth pupae from parasitoids. Arthropod-Plant Interactions, 7, 445–453.

Mackauer, M. & Sequeira, R. (1993) Patterns of development in insect parasites. Parasites and pathogens of insects, 1–23.

Miles, C.I., Chen, W.P., Adamo, S.A., Kester, K.M., & Miller, D.W. (2023) Manduca sexta caterpillars parasitized by the wasp Cotesia congregata stop chewing despite an intact motor system. J Exp Biol, 226.

Molleman, F., Sokół-Łętowska, A., Mallick, S., Prinzing, A., & Walczak, U. (2025) Adaptation to leaf traits of individual trees in a forest appears rare in caterpillars. Ecology and Evolution, 15, e71038.

Moore, M.E., Kester, K.M., & Kingsolver, J.G. (2020) Rearing temperature and parasitoid load determine host and parasitoid performance in Manduca sexta and Cotesia congregata. Ecological Entomology, 45, 79–89.

Price, P.W., Bouton, C.E., Gross, P., McPheron, B.A., Thompson, J.N., & Weis, A.E. (1980) Interactions among three trophic levels: influence of plants on interactions between insect herbivores and natural enemies. Annual Review of Ecology, Evolution, and Systematics, 11, 41–65.

R_Core_Team. (2025) R: A language and environment for statistical computing. R Foundation for Statistical Computing.

Razowski, J. (1983). Motyle (Lepidoptera) Polski. Część VI—Olethreutinae: Olethreutidii. In Monografie Fauny Polski;, Vol. 13. PWN, Warsaw-Krakow, Poland.

Razowski, J. (1987). Motyle (Lepidoptera) Polski. Część VII—Uzupełnienia i Eucosmini. In Monografie Fauny Polski, Vol. 15. PWN, Warszawa-Kraków, Poland.

Remmel, T., Davison, J., & Tammaru, T. (2011) Quantifying predation on folivorous insect larvae: the perspective of life-history evolution. Biological Journal of the Linnean Society, 104, 1–18.

Tammaru, T. & Esperk, T. (2007) Growth allometry of immature insects: larvae do not grow exponentially. Functional Ecology, 1099–1105.

Wheeler, J. (2023) Moth Dissection UK.

