## Appendix 1 growth curves for "Parasitoids reduce growth rate in oak-feeding caterpillars"

**Supplement 1:** **Growth curves of caterpillars**

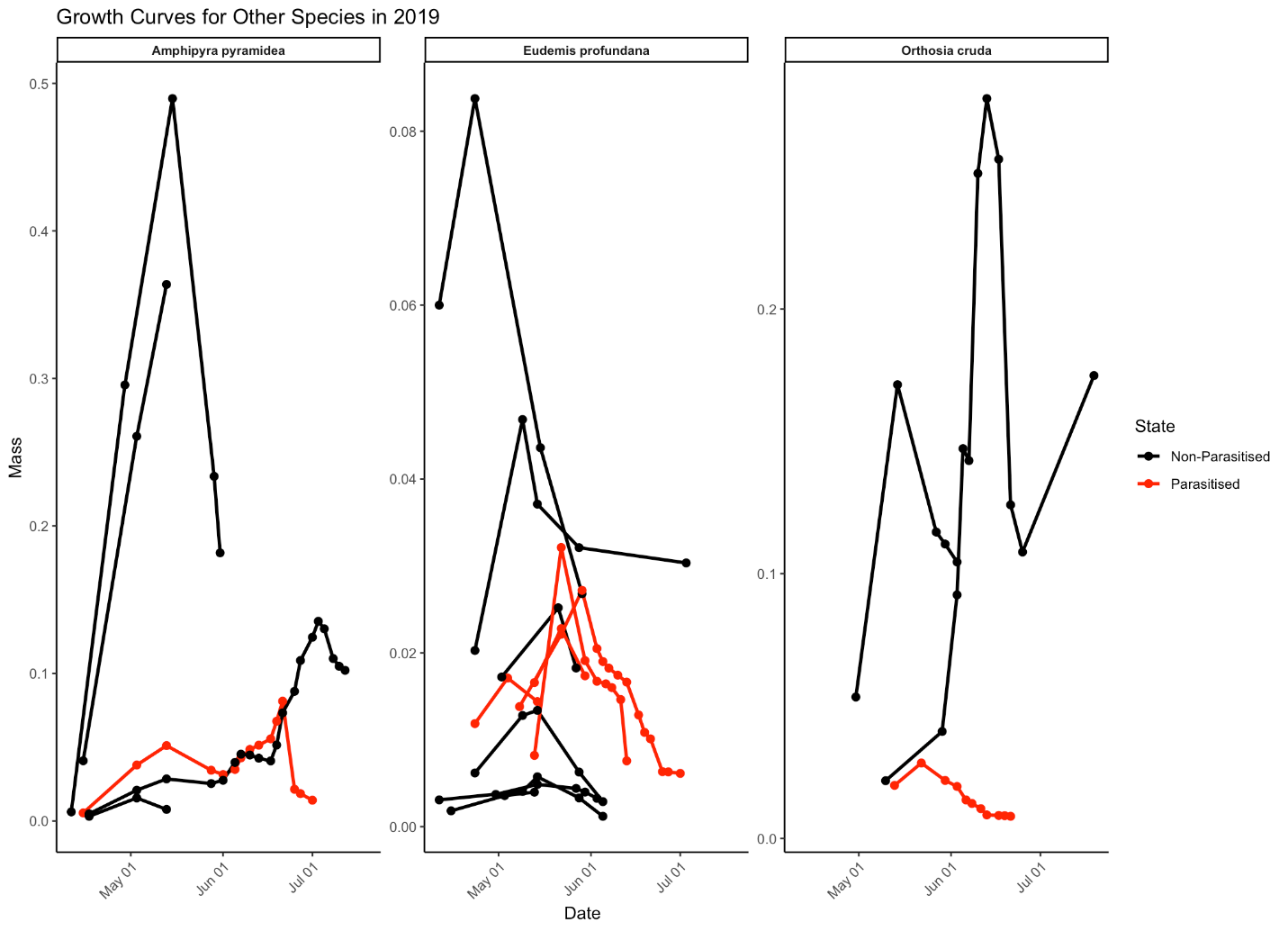

Figure S1.1. Growth trajectories of caterpillars collected in 2019, with mass changes over time for both parasitised (red data points) and non-parasitised individuals (black data points) where consecutive measurements are connected with a straight line. This excludes *Operopthera brumata* which are shown in the next graph.

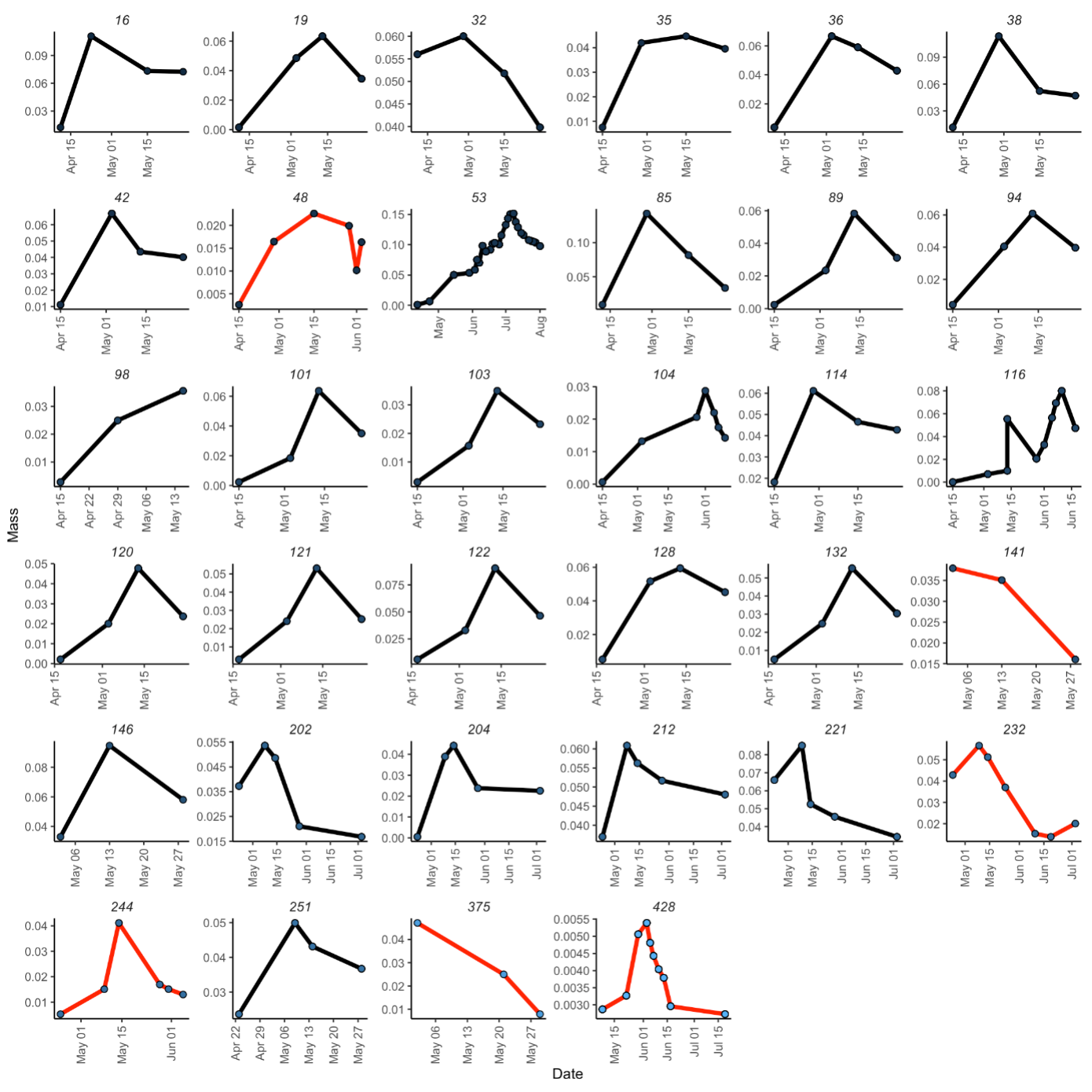

Figure S1.2: Growth curves for the *Operophtera brumata* individuals reared in 2019, showing mass changes over time for both parasitised (red lines) and non-parasitised (black lines) individuals. The graph includes 34 specimens, with 6 parasitised and 28 non-parasitised individuals. The y-axis represents mass, while the x-axis represents the date. Black lines indicate non-parasitised specimens, and red lines indicate parasitised specimens. The highest mass value for parasitised specimens is approximately 0.06 g, while the highest mass value for non-parasitised specimens reaches around 0.15 g. This visualization highlights the impact of parasitism on the growth patterns of *Operophtera brumata* caterpillars, showcasing noticeable differences in mass changes between parasitised and non-parasitised states.

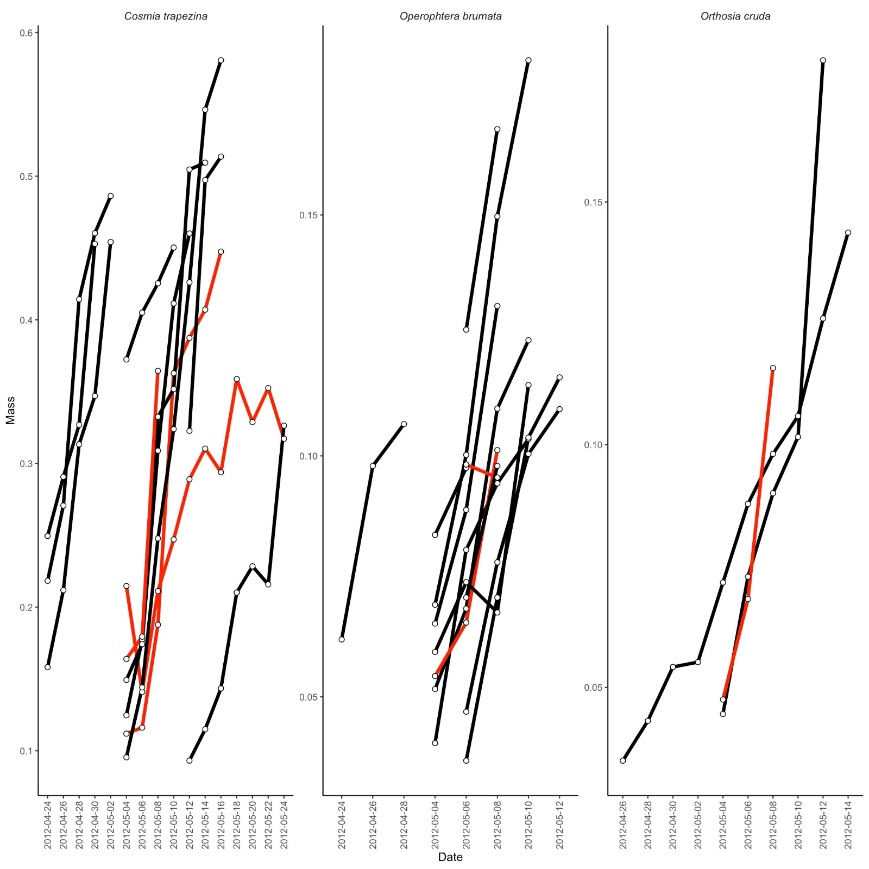

Figure S1.3: Growth curves for the individuals reared in 2012. Black lines indicate non-parasitised specimens, and red lines indicate parasitised specimens. Caterpillar body mass (mg) was inferred from their length.
